# Genomic Selection for End-Use Quality and Processing Traits in Soft White Winter Wheat Breeding Program with Machine and Deep Learning Models

**DOI:** 10.1101/2021.05.24.445513

**Authors:** Karansher S. Sandhu, Meriem Aoun, Craig Morris, Arron H. Carter

## Abstract

Breeding for grain yield, biotic and abiotic stress resistance, and end-use quality are important goals of wheat breeding programs. Screening for end-use quality traits is usually secondary to grain yield due to high labor needs, cost of testing, and large seed requirements for phenotyping. Hence, testing is delayed until later stages in the breeding program. Delayed phenotyping results in advancement of inferior end-use quality lines into the program. Genomic selection provides an alternative to predict performance using genome-wide markers. Due to large datasets in breeding programs, we explored the potential of the machine and deep learning models to predict fourteen end-use quality traits in a winter wheat breeding program. The population used consisted of 666 wheat genotypes screened for five years (2015-19) at two locations (Pullman and Lind, WA, USA). Nine different models, including two machine learning (random forest and support vector machine) and two deep learning models (convolutional neural network and multilayer perceptron), were explored for cross-validation, forward, and across locations predictions. The prediction accuracies for different traits varied from 0.45-0.81, 0.29-0.55, and 0.27-0.50 under cross-validation, forward, and across location predictions. In general, forward prediction accuracies kept increasing over time due to increments in training data size and was more evident for machine and deep learning models. Deep learning models performed superior over the traditional ridge regression best linear unbiased prediction (RRBLUP) and Bayesian models under all prediction scenarios. The high accuracy observed for end-use quality traits in this study support predicting them in early generations, leading to the advancement of superior genotypes to more extensive grain yield trailing. Furthermore, the superior performance of machine and deep learning models strengthen the idea to include them in large scale breeding programs for predicting complex traits.

## Introduction

Wheat (*Triticum aestivum* L.) breeding programs mainly focus on improving grain yield, biotic and abiotic stress tolerance, and end-use quality traits. Hexaploid wheat is classified into hard and soft wheat classes based on kernel texture, milling quality, protein strength, and water absorption (Souza et al. 2002). Soft wheat flour has lower damaged starch, gluten strengthen, and non-starch polysaccharides leading to less water absorption. In contrast, hard wheat has higher damaged starch, gluten strengthen, and non-starch polysaccharides causing higher water absorption (Kiszonas et al. 2013). Hard wheat dough is mainly used for pan type, leavened bread, flatbread, and noodles, whereas soft wheat dough is primarily used for cookies, cakes, and confectionery products (Bhave and Morris 2008; Kiszonas et al. 2013). Washington state was ranked fourth in the nation’s wheat production in 2020. About 80% of wheat grown in eastern Washington is soft white wheat (SWW), one of the six class grown in the USA. SWW is the smallest wheat class and is consistently in demand from overseas markets owing to its end-use quality attributes. More than 85% of the SWW produced in the Pacific Northwest (PNW) region is exported to markets in countries like Japan, Korea, the Philippines, and Indonesia.

End-use quality and processing traits are the combinations of various predefined parameters. Multiple attributes are measured from milling traits, baking parameters, grain characteristics, and flour parameters to assess product quality (Guzman et al. 2016). Milling traits are measured to extract flour and break flour percentage as flour yield and break flour yield (Morris et al. 2009). In general, soft wheat has a higher break flour yield than hard wheat. Thermogravimetric ovens are used for calculating the flour ash. Lower flour ash is recommended as higher amounts of minerals in ash reduces the functionality of most dough and batters (Morris et al. 2009). The milling score is estimated using flour yield, break flour yield, and flour ash content and is described in the Material and Methods section. The sugar snap cookie test is a must for SWW testing to me expectations of product performance from overseas markets. Baking of cooking is performed for lines within the breeding program, and SWW lines having cookie diameter above 9.3 cm is preferred (Kiszonas et al. 2015).

Grain characteristics commonly measured in SWW include kernel hardness, kernel size, kernel weight, test weight, and grain protein content. Kernel weight, kernel size, and kernel texture (hardness) are measured with a single kernel characterization system (SKCS). Lower values from the SKCS demonstrate softness; thus, SKCS values are negatively correlated with break flour yield. However, the two measures of kernel texture are not entirely correlated because SKCS includes only kernel resistance while break flour yield includes particle size and grain structure (Campbell et al. 2007). Grain and flour protein content plays a critical role in confectionery products from SWW. High gluten strength or viscoelastic strength is required for bread baking, whereas confectionary products require less gluten and water absorption. Gluten strength, and water absorption capacity, is measured using sodium dodecyl sulfate sedimentation and water solvent retention capacity tests. Lower water absorption in SWW aid in better cookie spread. Moreover, a flour swelling volume test is conducted to determine the amount of amylose and amylopectin components in the grain starch. Larger amylopectin content leads to higher flour swelling volume value, resulting in waxy starches required for certain Asian-style noodles (Kiszonas et al. 2013; Guzman et al. 2016).

Major genes influencing end-use quality traits are typically already fixed in most breeding programs, especially in different market classes. Until now, marker-assisted selection has been used for major genes controlling end-use quality, namely, low molecular weight glutenins, high molecular weight glutenins, granule bound starch synthase 1 (amylose composition) and puroindolines (kernel hardness) (Gale 2005; Kiszonas et al. 2013). Usage of these molecular markers only aid in differentiating different wheat classes earlier in the breeding program; however, they do not provide the complete profile of different end-use quality traits. Previous linkage mapping and genome-wide association studies in SWW have shown that a large number of small effect QTLs control most end-use quality traits in addition to the already fixed genes (Carter et al. 2012; Jernigan et al. 2018). Similarly, 299 small effects QTLs were identified using multi-locus genome-wide association studies for nine end-use quality traits in hard wheat (Yang et al. 2020). Kristensen et al. (2018) were unable to identify significant QTLs for Zeleny sedimentation, grain protein content, test weight, thousand kernel weight, and falling number in wheat and suggested genomic selection as the best alternative for predicting quantitative traits.

Genomic selection (GS) opens up the potential for selecting improved end-use quality lines due to the small effect of these loci, limited seed availability earlier in the breeding pipeline for conducting tests, and time constraints in winter wheat for sowing the new cycle (Crossa et al. 2017). GS uses the genotypic and phenotypic data from previous breeding lines or populations to train predictive statistical models. These trained models are subsequently used to predict the genomic estimated breeding estimated values (GEBVs) of genotyped lines (Meuwissen et al. 2001). GS has shown the potential to enhance genetic gain by reducing the generation advance time and improving selection accuracy (Battenfield et al. 2016; Juliana et al. 2019; Sandhu et al. 2021b). This is especially important for winter wheat end-use quality traits, as phenotyping requires more than three months and data from the quality lab is often not available between harvest and the time planting occurs. This ultimately results in either the increase of one year in the breeding cycle or passage of undesirable lines into the next growing season. Furthermore, phenotyping requires a large amount of seed and is costly, so large-scale testing is often not conducted until later generations. Currently, the cost of genotyping 10,000 lines with high density genotyping by sequencing is equivalent to phenotyping 200 lines for end-use quality and processing traits (Guzman et al. 2016). GS is the best technique for breeding end-use quality traits after considering time, cost, and seed amount.

Genomic selection has been primarily explored in several hard wheat end-use quality trait studies using the traditional genomic best linear biased prediction (GBLUP), Bayes A, Bayes B, Bayes C, and Bayes Cpi, showing mixed results, where one model performed best for one trait and not for another (Heffner et al. 2011a, b). Machine and deep learning models have opened up an entirely new platform for plant breeders and exploring them in the breeding program could accelerate the pace of genetic gain. Deep learning models have shown higher prediction accuracies for different complex traits in wheat (Sandhu et al. 2021a), rice (*Oryzae sativa* L.; Chu and Yu 2020), soybean (*Glycine max* L.; Liu et al. 2019), and maize (*Zea mays* L.; Khaki and Wang 2019). Sandhu et al. (2021a) have shown that two deep learning models, namely, convolutional neural network (CNN) and multilayer perceptron (MLP), gave 1-5% higher prediction accuracy compared to BLUP based models. Ma et al. (2018) and Montesinos-López et al. (2019) also obtained similar results to predict quantitative traits in wheat and suggested that deep learning models should be explored due to their better prediction accuracies. To the best of our literature search, this is the first study exploring the potential of the deep learning models for predicting the end-use quality traits in wheat.

This study explored the potential of GS using multi-environment data from 2015-19 for end-use quality traits in a soft white winter wheat breeding program. We explored nine different BLUP based models, Bayesian models, and machine and deep learning models to predict the fourteen different end-use quality traits. The main objectives of this include, 1) Optimization of the machine and deep learning models for predicting end-use quality traits, 2) Comparison of prediction ability of nine different GS models to predict fourteen different end-use quality traits using cross-validation approaches, and 3) Assess the potential of GS for forward prediction and across location predictions using previous years training data in the breeding program.

## Materials and Methods

### Germplasm

A total of 666 soft white winter wheat lines were evaluated for five years at two locations, namely, Pullman and Lind, WA, USA, from 2015-19. These 666 genotypes consist of F_4:5_ derived lines, double haploid lines, lines in preliminary and advanced yield trials screened as a part of the Washington State University winter wheat breeding program. F_4:5_ derived lines and double haploid lines were screened for the agronomic and disease resistance traits, and the superior genotypes were tested for the end-use quality. Lines in preliminary and advanced yield trials were selected for superior yield, and those lines were later advanced for end-use quality traits phenotyping. Some genotypes were replicated at a single location per year, whereas others were un-replicated, creating an unbalanced dataset.

### Phenotyping

Fourteen different end-use quality and processing traits were measured, and data were obtained from the USDA-ARS Western Wheat Quality Laboratory, Pullman, WA. All these traits were measured following the guidelines of the American Association of Cereal Chemists International (AACCI 2008). These fourteen traits were divided into four categories: milling traits, baking parameters, grain characteristics, and flour parameters. The complete summary of each trait, number of observations, mean, standard error, and heritability is provided in **Table 1 & Table 2**.

**Table 1.** Total number of lines screened across each year at two locations in Washington and phenotyped for enduse quality traits.

| Location | Year | Lines screened for quality |
| --- | --- | --- |
| Lind | 2015 | 122 |
|  | 2016 | 114 |
|  | 2017 | 115 |
|  | 2018 | 71 |
|  | 2019 | 106 |
| Pullman | 2015 | 183 |
|  | 2016 | 128 |
|  | 2017 | 181 |
|  | 2018 | 137 |
|  | 2019 | 178 |
| Total |  | 1335 |

**Table 2.** Summary of the fourteen end-use quality traits evaluated for genomic selection analysis using nine different prediction models.

| Trait | Abbreviation | Units | Number of genotypes | Mean | Min | Max | S.E. | H<sup>2</sup> |
| --- | --- | --- | --- | --- | --- | --- | --- | --- |
| **Milling traits** |  |  |  |  |  |  |  |  |
| FYELD | Flour yield | percent | 666 | 69.9 | 58.0 | 75.8 | 0.09 | 0.91 |
| BKYELD | Break flour yield | percent | 666 | 48.1 | 33.9 | 56.6 | 0.14 | 0.93 |
| MSCOR | Milling score | unitless | 646 | 85.6 | 69.1 | 98.8 | 0.10 | 0.81 |
| **Grain characteristics** |  |  |  |  |  |  |  |  |
| TWT | Test weight | Kg/hL | 666 | 61.8 | 54.6 | 65.9 | 0.06 | 0.92 |
| GPC | Grain protein content | percent | 666 | 10.73 | 7.2 | 14.8 | 0.05 | 0.56 |
| KHRD | Kernel hardness | unitless | 666 | 23.0 | -10.2 | 52.4 | 0.4 | 0.93 |
| KWT | Kernel weight | mg | 666 | 39.3 | 26.5 | 54.6 | 0.17 | 0.86 |
| KSIZE | Kernel size | mm | 666 | 2.76 | 2.3 | 3.3 | 0.005 | 0.83 |
| **Baking parameters** |  |  |  |  |  |  |  |  |
| CODI | Cookie diameter | cm | 622 | 9.2 | 7.8 | 10.0 | 0.008 | 0.89 |
| **Flour parameters** |  |  |  |  |  |  |  |  |
| FPROT | Flour protein | percent | 666 | 8.93 | 6.3 | 13.0 | 0.04 | 0.57 |
| FASH | Flour ash | percent | 646 | 0.39 | 0.21 | 0.54 | 0.001 | 0.88 |
| FSV | Flour swelling volume | mL/g | 665 | 19.06 | 14.0 | 26.3 | 0.05 | 0.63 |
| FSDS | Flour SDS sedimentation | g/mL | 666 | 10.1 | 3.5 | 18.3 | 0.09 | 0.92 |
| FSRW | Water solvent retention capacity in water | percent | 666 | 54.18 | 43.4 | 72.6 | 0.09 | 0.85 |

S.E. is standard error H<sup>2</sup> is broad sense heritability

Grain characteristics, namely kernel size (KSIZE), kernel weight (KWT), and kernel hardness (KHRD) were determined using 200 seeds/sample with a SKCS 4100 (Perten Instruments, Springfield, IL, USA) (AACC Approved Method 55-31.01). Grain protein content (GPC) was measured using a NIR analyzer (Perten Elmer, Sweden) (AACC Approved Method 39-10.01). Test weight (TWT) was obtained as weight/volume following AACC Approved Method 55-10.01.

Three milling traits, namely flour yield (FYELD), break flour yield (BKYELD), and milling score (MSCOR) were obtained using a Quadrumat senior experimental mill (Brabender, South Hackensack, NJ, USA). FYELD was determined as a ratio of total flour weight (mids + break flour) to the initial sample weight using a single pass through the Quadrumat break roll unit. BKYELD was estimated as the percent of milled product passing through a 94-mesh* screen per unit grain weight. Flour ash (FASH) was obtained using the AACC Approved Method 08-01.01. MSCOR was calculated using the formula: MSCOR= (100–(0.5(16–13.0 + (80–FYELD) + 50 (FASH–0.30))) × 1.274) –21.602, showing that this trait is a function of FYELD and FASH content. To evaluate baking parameters, cookie diameter (CODI) was measured using AACC Approved Method 10-52.02.

Four different flour parameters, namely, flour protein (FPROT), water solvent retention capacity in water (FSRW), flour swelling volume (FSV) and flour sodium dodecyl sulfate sedimentation (FSDS) were measured from the extracted flour. FPROT was measured following the AACC Approved Method 39-11.01. FSRW measures the water retention capacity of gluten, gliadins, starch, and arabinoxylans using the AACC Approved Method 56-11.02. The FSDS test was used to measure strength of gluten by following the AACC Approved Method 56-60.01. The FSV test assesses starch composition following the AACC Approved Method 56-21.01.

### Statistical analysis

Due to the unbalanced nature of the dataset, adjusted means were calculated using residuals obtained using the lme4 R package for within environment analysis. The model equation is represented as

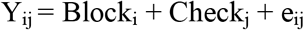

Where Y_ij_ is the raw phenotype; Checkj is the effect of replicated check cultivar; Blocki corresponds to the fixed block effect; and e_ij_ is the residuals (Bates et al. 2015).

Adjusted means across the environments were calculated following the method implemented in Sandhu et al. (2021c) and is as follows

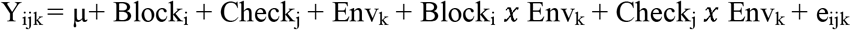

Where Y_ijk_ is the raw phenotype value; Block_i_, Check_j_, and Env_k_ are the fixed effect of ith block, jth check, and kth environment; and e_ijk_ is the residuals.

Best linear unbiased predictors (BLUPs) for individuals and across environments were used to obtain the variance components for estimating broad sense heritability. The equation for heritability used was

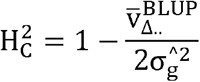

Where 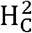 is the Cullis heritability; 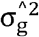 is genotypic variance; and 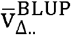 is the mean-variance of BLUPs (Cullis et al. 2006).

### Genotyping

The whole population was genotyped using GBS through the North Carolina State University (NCSU) Genomics Sciences Laboratory, Raleigh, NC, using the restriction enzymes *Pst*I and *Msp*I (Poland et al. 2012). LGC Biosearch Technologies Oktopure^™^ robotic platform with sbeadex^™^ magnetic microparticle reagent kits were used to extract the DNA from the leaves of ten-day-old seedlings. Thermo Fisher (Waltham, MA) Quant-It^™^ PicoGreen^™^ assays were used to quantify the DNA, and the samples were normalized to 20 ng/μL. Restriction enzymes *Pst*I and *Msp*I were used for sample fragmentation, and the digested samples were ligated with barcode adapters using T4 ligase. The pooled samples were amplified using PCR, following Poland et al. (2012), and sequencing was performed at NCSU Genomics Sciences Laboratory. Burrows-Wheeler Aligner (BWA) 0.7.17 was used to align the sequences to the Chinese Spring (IWGSC) RefSeq v1.0 reference genome (Appels et al. 2018). Tassel v5 and Beagle were used for SNP discovery, calling, and imputation (Bradbury et al. 2007). Quality filtering pipeline was implemented in R software to remove markers with minor allele frequency less than 5%, markers missing more than 20% data, and heterozygosity more than 15%. After the complete filtering pipeline, 40,518 SNPs remained and used for population structure and genomic prediction.

### Genomic selection models

We explored the performance of five parametric and four non-parametric models for all fourteen traits evaluated in this study. Parametric models used were RRBLUP, Bayes B, Bayes A, Bayes Lasso, and Bayes C. Non-parametric models included two machine and two deep learning models. The complete information for all those models and optimization process is provided as follows:

### Ridge regression best linear unbiased prediction (RRBLUP)

RRBLUP was included here as the benchmark for comparing its performance with other models due to frequent use in wheat breeding and ease of implementation. The model assumes that all markers contribute to the trait and has a constant effect variance. Marker effects and variance patterns are estimated using the restricted estimated maximum likelihood (REML) function based on phenotypic and marker data (Endelman 2011). The RRBLUP model was implemented with the R package rrBLUP using the *mixed.solve* function. The model can be represented as

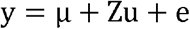

Where μ is the overall mean; y is the vector of adjusted means; u is a vector with normally distributed random marker effects with constant variance as u ~ N(0, I***σ***^2^_u_); Z is an N x M matrix of markers; and e is the residual error distributed as e ~ N(0, I***σ***^2^_e_). The solution for mixed equation can be written as

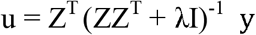

Where u, Z and y are explained above; I is an identity matrix and λ is represented as λ = ***σ***^2^_e_ / σ^2^_u_ and is the ridge regression parameter (Endelman 2011).

### Bayesian models

We implemented four different Bayesian models, namely, Bayes Lasso, Bayes A, Bayes B, and Bayes C. All these models assume different prior distributions for estimating marker effects and variances. Bayes A applies the inverted chi-squared probability distribution for estimating marker variances. Bayes B provides a more realistic scenario for breeding, assuming that all markers do not contribute to total genetic variation. It applies a mixture of prior distribution with a high probability mass at zero, and others follow the Gaussian distribution. Bayes C and Bayes Lasso follow the mixture of the prior distribution (point mass at zero with scaled-t distribution) and long-tail student t distribution (Pérez and Campos 2014). All the Bayesian models were implemented using the BGLR R package using the model equation

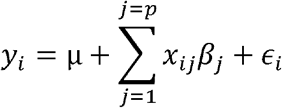

Where μ, *y_i_, x_ij_*, and *ϵ_i_* are defined above; and *β_j_* is the jth marker effect. Each Bayesian model used in this study has separate conditional prior distribution. Analysis was performed with 30,000 Monte Carlo Markov chain iterations with 10,000 burn-in iterations (Pérez and Campos 2014).

### Random forests (RF)

RF involves building a large collection of identical distributed trees and averages from the trees for final prediction. Different bootstrap samples are performed over the training set to identify the best feature subsets for splitting the tree nodes. The main criteria for splitting at the node include lowering the loss function during each bootstrapped sample (Shah et al. 2019). Model equation is represented as

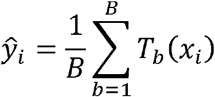

Where *ŷ_i_* is the predicted value of the individual with genotype *x_i_*; *T* is the total number of trees; and *B* is the number of bootstrap samples. The main steps involved in model functioning includes

1. Bootstrap sampling (b = (1,…, B)) to select plants with replacement from the training set, and an individual plant can appear once or several time during the sampling
2. Best set of features (SNPj, j = (1,…, J) were selected to minimize the mean square error (MSE) using the max feature function in the random forest regression library.
3. Splitting is performed at each node of the tree using the SNPj genotype to lower the MSE.
4. The above steps are repeated until a maximum depth is reached or a minimum node. The final predicted value of an individual of genotype *x_i_* is the average of the values from the set of trees in the forest.

The important hyperparameter model training include the depth of the trees, the importance of each feature, the number of features sampled for each iteration, and the number of trees. Randomized grid search cross-validation was used for hyperparameter optimization. The combination of hyperparameters that were tried included max depth (40, 60, 80, 100), max features (auto, sqrt), and number of trees (200, 300, 500, 1000) (Hastie et al. 2009). The Scikit learn, and random forest regression libraries in Python 3.7 were used for analysis (Gulli and Pal 2017).

### Support vector machine (SVM)

SVM uses the non-linear kernel for mapping the predictor space to high dimensional feature space for studying the relationship between marker genotype and phenotypes. The model equation is represented as

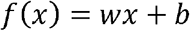

Where *f*(*x*) is learning function; *b* is the constant, reflecting the maximum allowed bias; *w* is the unknown weight; and x is the marker set. The learning function is mapped by minimizing the loss function as

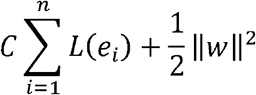

Where *C* is a positive regularization parameter; ║*w*║^2^ represents model complexity, *e_i_* = y - *f*(*x*) is the associated error with the *i^th^* training data point, and *L* is the loss function (Smola and Scholkopf 2004).

### Multilayer perceptron (MLP)

MLP is the feed-forward deep learning model that uses three layers, namely, input, hidden, and output, for mapping the relationship. These layers are connected by a dense network of neurons, where each neuron has its characteristic weight. MLP uses the combination of neurons, activation function, learning rate, hidden layers, and regularization for predicting the phenotypes. Input layer corresponds to SNP genotypes while neurons connect multiple hidden layer with associated strength (weight). The output of the *i^th^* hidden layer is represented as

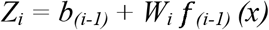

Where *Z_i_* is the output from the *i^th^* hidden layer; b_0_ is the bias for estimating neurons weight; *f_(i-1)_* represents the activation function; and *W_i_* is the weight associated with the neurons, and this process is repeated until the output layer.

Keras function’s grid search cross-validation and internal capabilities were used for optimizing the hyperparameters. Hyperparameters giving the lowest MSE were identified and used for output prediction (Cho and Hegde 2019). Regularization, dropout, and early stopping were applied to control overfitting. Furthermore, information about hyperparameter optimization and deep learning models is referred to in Sandhu et al. (2021a, c).

### Convolutional neural network (CNN)

CNN is a special case of deep learning model that accounts for the specific pattern present between the input features. Information about the CNN model, its implementation, and hyperparameter optimization are referred to in previous publications (Sandhu et al. 2021a, d). A combination of input, convolutional, pooling, dense, flatten, dropout, and output layers were applied for the prediction. Like MLP, hyperparameter was optimized using grid search cross-validation to select filters, activation function, solver, batch size, and learning rate. Regularization, dropout, and early stopping were applied to control overfitting. All the deep learning algorithms were implemented using Scikit learn and Keras libraries (Pedregosa et al. 2011; Srivastava et al. 2014).

### Prediction accuracy and cross-validation scheme

Prediction accuracy was evaluated using five-fold cross-validation where 20% of the data was used for testing and the remaining 80% for training within each environment. One hundred replications were performed for assessing each model’s performance. One replicate consisted of five iterations where data is split into five different groups. Prediction accuracy was reported as the Pearson correlation coefficient between the true (observed phenotype) and GEBVs. Separate analysis was performed for both locations using a cross-validation approach to assess the model’s performance.

Independent predictions or forward predictions were performed by training the model on previous year data and predicting future environments (i.e., 2015 data from Lind was used to predict 2016; 2015 and 2016 data predicts 2017, and so on for both locations). In the end, we tried to predict the 2019 environment of both locations by using the whole data set from the other location (i.e., 2015-19 data from Lind was used to predict 2019 in Pullman). Forward prediction represents real prediction scenarios in breeding programs where previous data are used to predict future environments. Due to computational burden, all the GS models were analyzed over the Kamiak high-performance cluster (https://hpc.wsu.edu/).

## Results

### Phenotypic data summary

**Table 1** provides the information about different lines screened for end-use quality traits across years at two locations. One thousand three hundred thirty-five lines were phenotypically screened for end-use quality traits across five years (2015-19) at two locations (**Table 1**). Overall, Pullman had more lines compared to Lind for each year. Summary statistics, including mean, minimum, maximum, standard error, and heritability are provided for all the fourteen end-use quality traits (**Table 2**). Broad sense heritability ranged from 0.56 to 0.93 for different traits. All the traits were highly heritable except GPC and FPROT (**Table 2**).

Significant positive and negative correlations were observed among different traits (**Figure 1**). Moderately high positive correlations were observed between FYELD and BKFYELD, KSIZE and KWT, GPC and FPROT, FSDS and FPROT, GPC and FSDS, and FSRW and KHRD (**Figure 1**). Similarly, moderately high negative correlations were seen between FASH and MSCOR, CODI and KHRD, GPC and FSV, FSDS and CODI, and CODI and FSRW (**Figure 1**). Most of the traits were not strongly correlated with each other, suggesting that a single quality trait cannot substitute others; hence, measurements from all of them are required for selection decisions.

**Figure 1:**
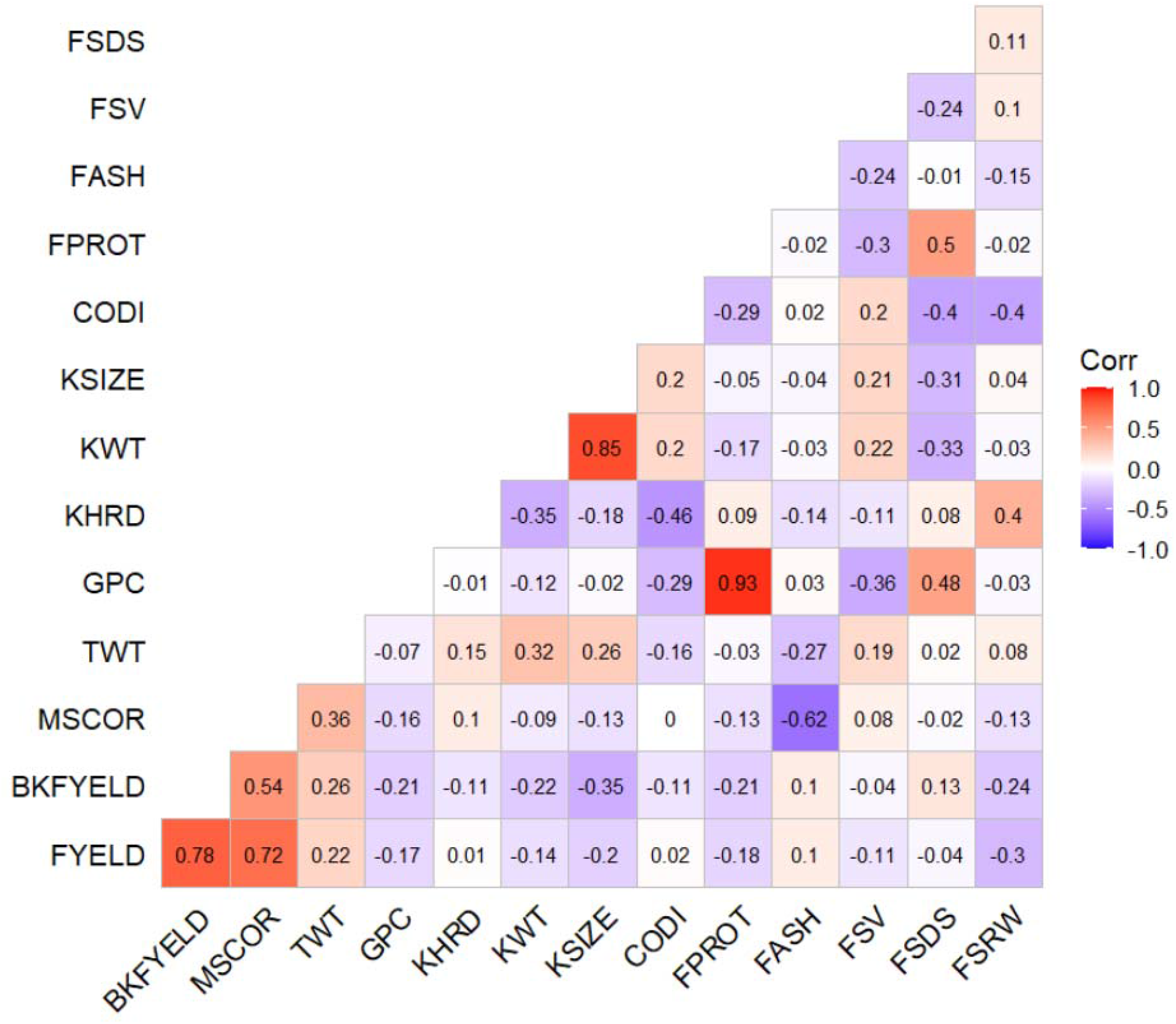
Phenotypic correlation between different end-use quality traits evaluated across two locations in Washington and five years using best linear unbiased predictors. All the abbreviation are previously abbreviated in the text and Table 2.

### Cross-validation genomic selection accuracy and model comparison

Complete datasets across the years from Pullman and Lind were used to predict the fourteen end-use quality traits using nine different models (**Table 3, Figure 2**). Five-fold cross-validation was performed to compare the results from the models at both locations. Prediction accuracy at Pullman varied from 0.52-0.81 for all traits with nine different GS models. The highest prediction accuracy was 0.81 for KWT and KSIZE with the RF and MLP model at Pullman (**Figure 2**). The lowest prediction accuracies were for GPC, FASH, FPROT, and FSRW at Pullman using different GS models (**Table 3**). The highest prediction accuracy for each trait is bolded for comparison with other models (**Table 3**). For the fourteen end-use quality traits evaluated in this study at Pullman, deep learning models, namely MLP and CNN, performed best for eight of the traits, demonstrating the potential to incorporate them into breeding programs (**Table 3**) for prediction. RF and SVM performed best for three and four traits out of the fourteen, while RRBLUP performed superior for only one trait (**Table 3 and Figure 2**).

**Figure 2.**
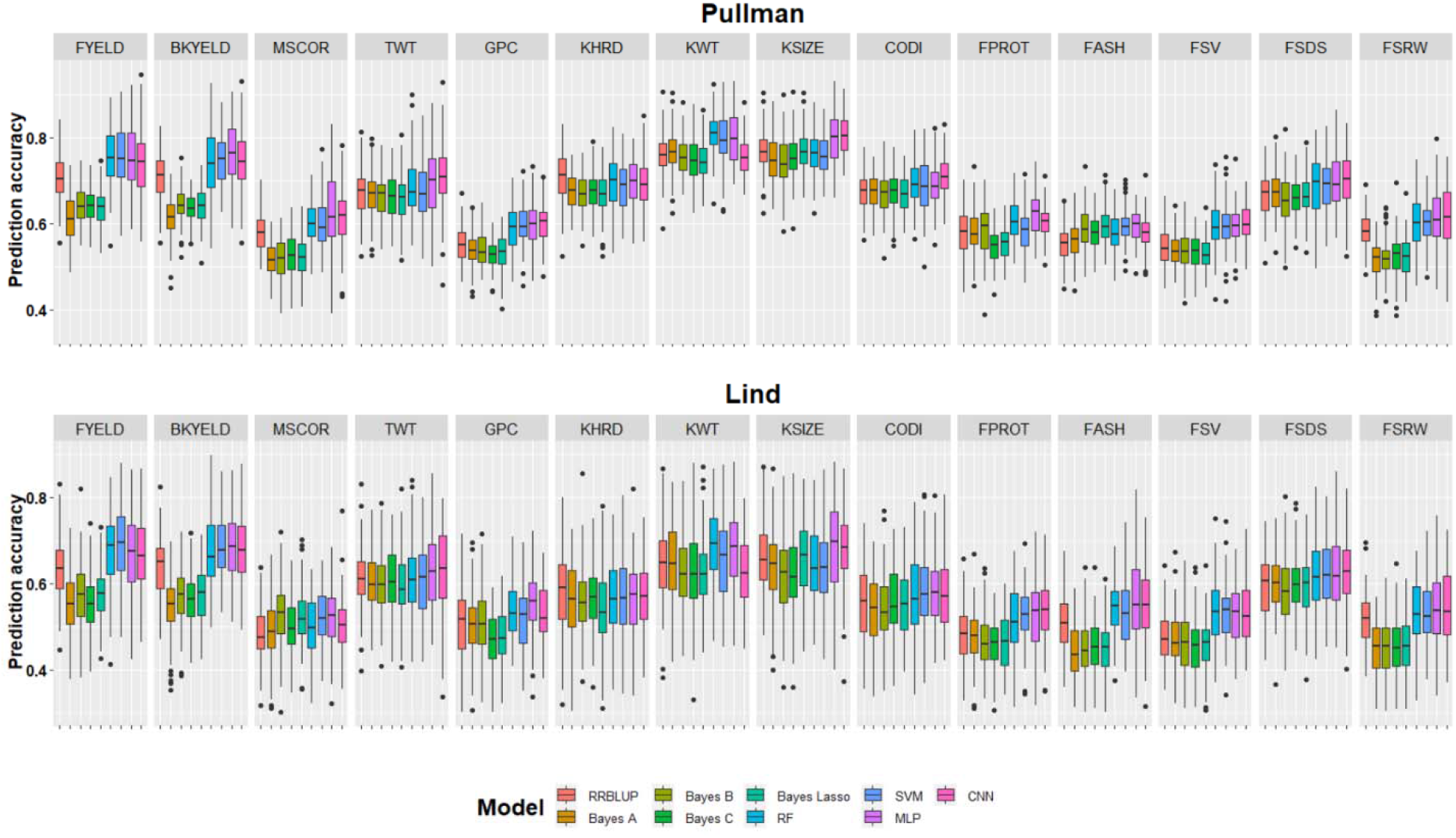
Genomic selection cross-validation prediction accuracies for fourteen end-use quality traits evaluated with nine different models. Results are provided separately for both locations and each trait is separated with facets.

**Table 3.** Genomic selection cross-validation prediction accuracies for the fourteen end-use quality traits evaluated with nine different models at two locations in Washington. The highest accuracy for each trait is bolded under different model scenarios.

| Location | Trait | RRBLUP | BayesA | Bayes B | Bayes C | Bayes Lasso | RF | SVM | MLP | CNN |
| --- | --- | --- | --- | --- | --- | --- | --- | --- | --- | --- |
| **Pullman** | FYELD | 0.71 | 0.61 | 0.64 | 0.64 | 0.63 | **0.76** | **0.76** | 0.75 | 0.74 |
|  | BKYELD | 0.70 | 0.62 | 0.64 | 0.64 | 0.64 | 0.75 | 0.75 | **0.76** | 0.75 |
|  | MSCOR | 0.58 | 0.52 | 0.52 | 0.53 | 0.52 | 0.60 | 0.60 | **0.63** | 0.61 |
|  | TWT | 0.67 | 0.67 | 0.66 | 0.66 | 0.66 | 0.68 | 0.67 | **0.70** | **0.70** |
|  | GPC | 0.55 | 0.54 | 0.54 | 0.53 | 0.53 | 0.59 | **0.60** | **0.60** | **0.60** |
|  | KHRD | **0.71** | 0.67 | 0.67 | 0.68 | 0.67 | 0.70 | 0.69 | 0.70 | 0.69 |
|  | KWT | 0.76 | 0.77 | 0.75 | 0.75 | 0.75 | **0.81** | 0.80 | 0.80 | 0.75 |
|  | KSIZE | 0.77 | 0.75 | 0.74 | 0.75 | 0.77 | 0.76 | 0.76 | 0.80 | **0.81** |
|  | CODI | 0.67 | 0.67 | 0.67 | 0.68 | 0.67 | 0.69 | 0.69 | 0.69 | **0.71** |
|  | FPROT | 0.58 | 0.58 | 0.58 | 0.55 | 0.55 | **0.61** | 0.58 | 0.62 | 0.60 |
|  | FASH | 0.55 | 0.56 | **0.59** | 0.58 | **0.59** | 0.58 | **0.59** | **0.59** | **0.59** |
|  | FSV | 0.55 | 0.54 | 0.53 | 0.53 | 0.53 | 0.59 | **0.60** | **0.60** | **0.60** |
|  | FSDS | 0.67 | 0.67 | 0.66 | 0.66 | 0.67 | 0.69 | 0.69 | **0.70** | **0.70** |
|  | FSRW | 0.58 | 0.52 | 0.52 | 0.52 | 0.52 | 0.60 | 0.60 | 0.61 | **0.62** |
| **Lind** | FYELD | 0.64 | 0.55 | 0.58 | 0.56 | 0.58 | 0.68 | **0.69** | 0.67 | 0.67 |
|  | BKYELD | 0.63 | 0.55 | 0.57 | 0.56 | 0.57 | 0.67 | 0.68 | **0.69** | **0.69** |
|  | MSCOR | 0.48 | 0.49 | **0.53** | 0.50 | 0.52 | 0.50 | 0.52 | 0.52 | 0.50 |
|  | TWT | 0.61 | 0.61 | 0.60 | 0.61 | 0.60 | 0.61 | 0.61 | 0.63 | **0.64** |
|  | GPC | 0.51 | 0.51 | 0.51 | 0.47 | 0.47 | 0.54 | 0.52 | **0.55** | 0.53 |
|  | KHRD | **0.58** | 0.56 | 0.56 | 0.57 | 0.54 | 0.56 | 0.57 | 0.57 | 0.57 |
|  | KWT | 0.65 | 0.65 | 0.63 | 0.63 | 0.63 | **0.70** | 0.66 | 0.69 | 0.63 |
|  | KSIZE | 0.66 | 0.64 | 0.62 | 0.63 | 0.66 | 0.64 | 0.64 | **0.69** | 0.68 |
|  | CODI | 0.56 | 0.54 | 0.54 | 0.56 | 0.55 | 0.57 | **0.58** | **0.58** | **0.58** |
|  | FPROT | 0.48 | 0.48 | 0.46 | 0.46 | 0.46 | 0.51 | 0.53 | 0.53 | **0.54** |
|  | FASH | 0.51 | 0.44 | 0.44 | 0.45 | 0.44 | 0.54 | 0.53 | **0.56** | 0.53 |
|  | FSV | 0.48 | 0.47 | 0.46 | 0.45 | 0.46 | **0.54** | **0.54** | 0.53 | 0.53 |
|  | FSDS | 0.59 | 0.60 | 0.59 | 0.60 | 0.59 | 0.62 | **0.63** | **0.63** | 0.62 |
|  | FSRW | 0.52 | 0.45 | 0.45 | 0.45 | 0.46 | 0.53 | 0.53 | **0.54** | **0.54** |
| **Average** |  | **0.61** | **0.58** | **0.58** | **0.58** | **0.58** | **0.63** | **0.63** | **0.64** | **0.63** |

All the abbreviation are previously abbreviated in the text and Table 2.

Prediction accuracies (0.454-0.70) within the Lind dataset were lower than Pullman for all traits (**Table 2**). Similar to Pullman, the highest cross-validation prediction accuracy (i.e. 0.70) was obtained for KWT at Lind. The lowest prediction accuracies were obtained for GPC, FPROT, and FSRW using the Bayesian models (**Table 3**). Machine and deep learning models performed superior for twelve out of the fourteen end-use quality traits (**Figure 2**). **Table 3** provides the average performance for all models, and we observed that machine and deep learning models performed superior to RRBLUP and all the Bayesian models. On average, machine and deep learning models performed 10% and 5%, superior to Bayesian and RRBLUP. Due to Bayesian model’s inferior performances and computational burden, they were not included for across location predictions (**Figure 5**).

**Figure 3.**
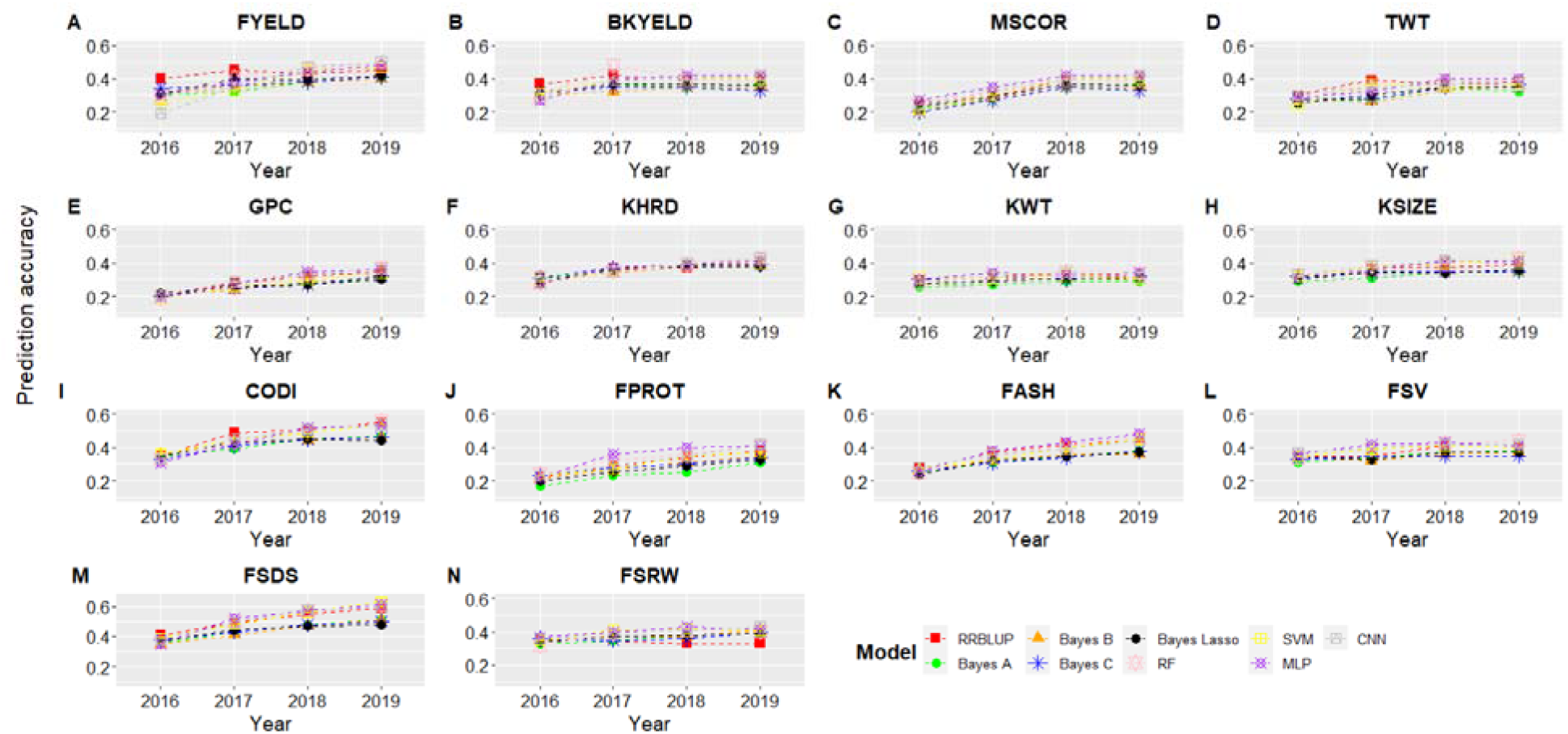
Genomic selection forward prediction accuracies for Pullman, WA, when all datasets from previous years were included to predict fourteen end-use quality traits using nine different models. The x-axis represents the year for which predictions were made using previous years as training set. All abbreviations are previously abbreviated in the text and Table 2.

**Figure 4.**
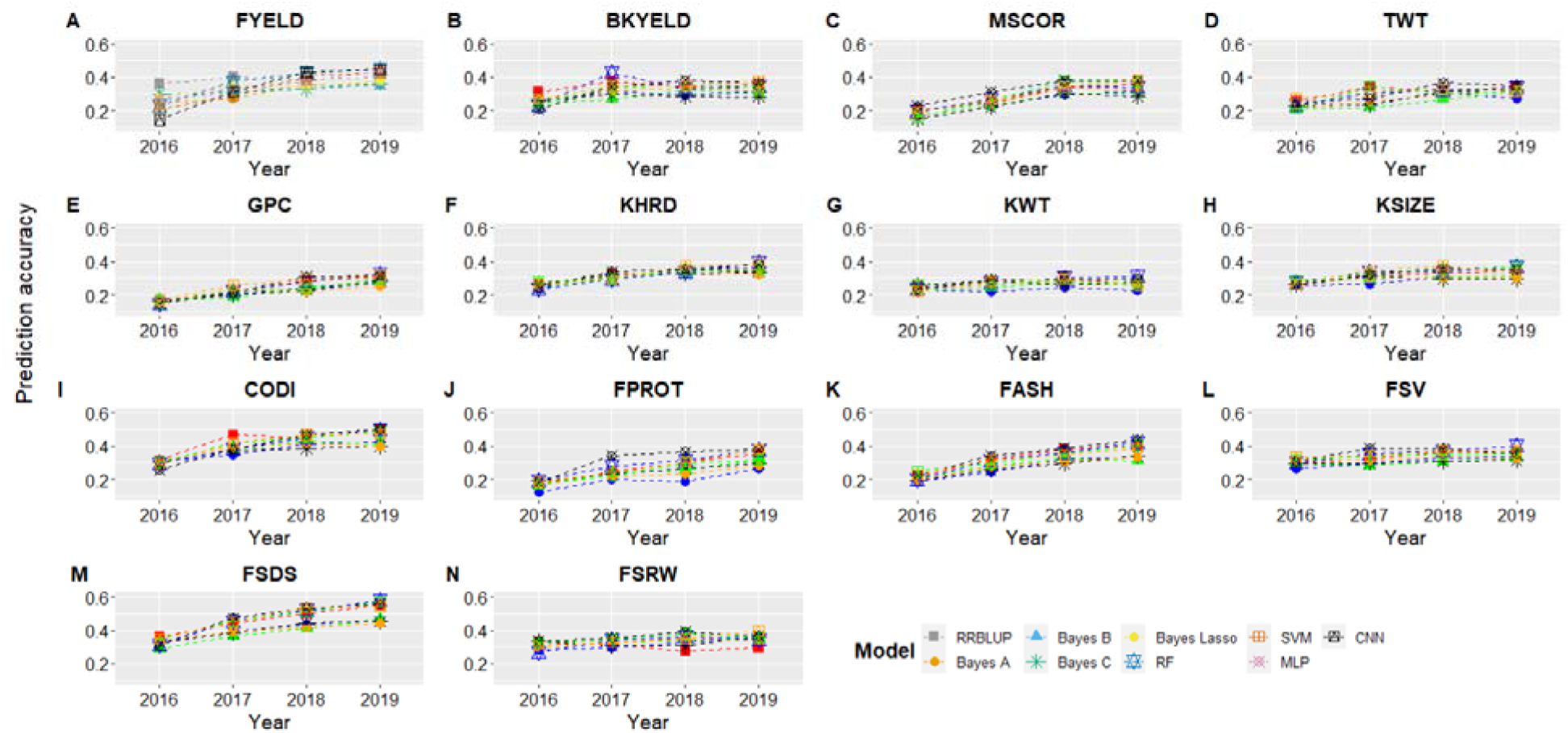
Genomic selection forward prediction accuracies for Lind, WA, when all datasets from previous years were included to predict fourteen end-use quality traits using nine different models. The x-axis represents the year for which predictions were made using previous years as the training set. All abbreviations are previously abbreviated in the text and Table 2.

**Figure 5.**
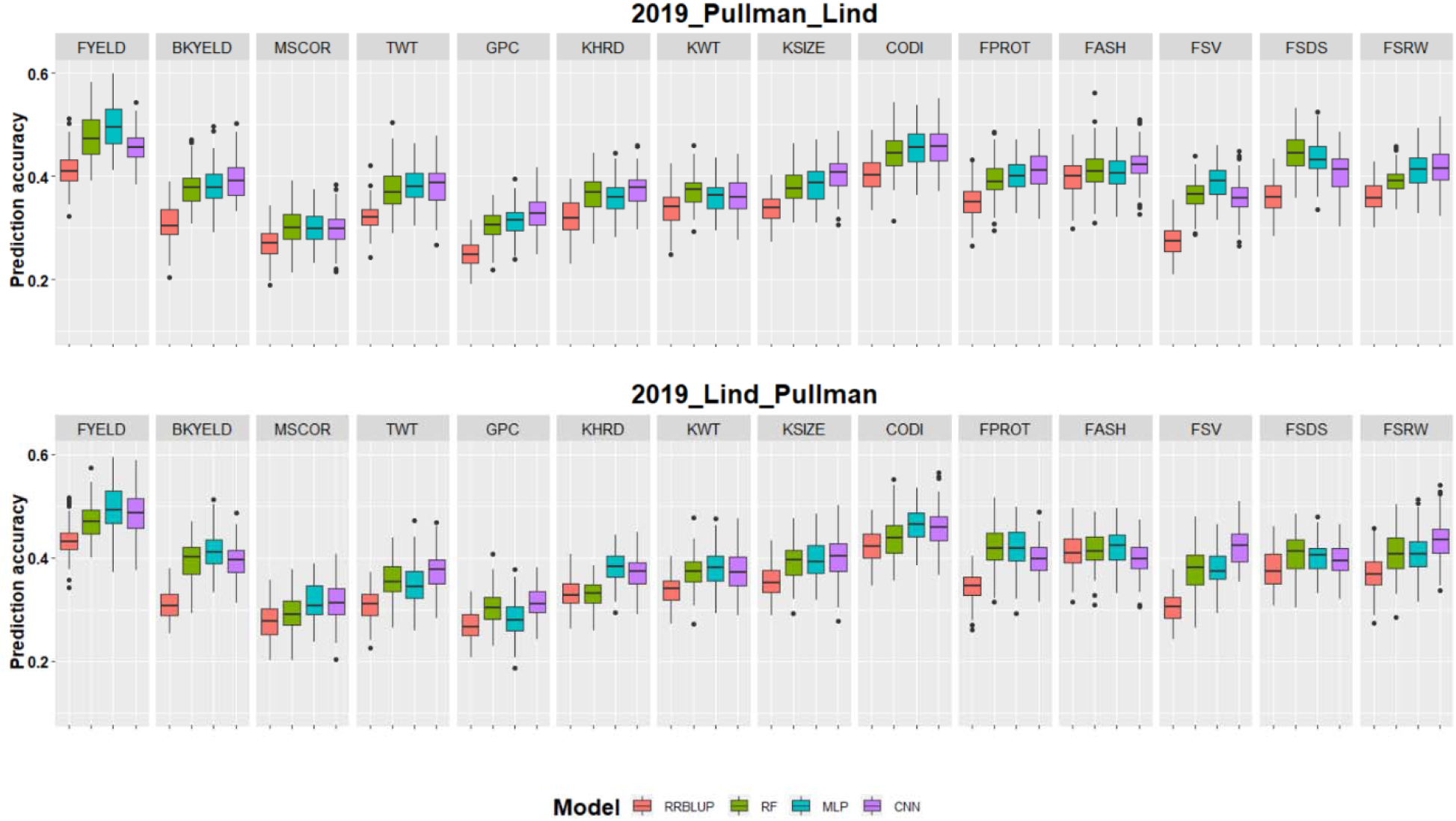
Genomic selection across environment prediction accuracies for fourteen end-use quality traits evaluated with four different models. 2019_Pullan_Lind denotes the scenario where 2019_Pullman was predicted using datasets from Lind as training set and vice versa for 2019_Lind_Pullman. Results are provided separately for both locations and each trait is separated with facets.

### Forward predictions

GS model predictions were assessed to reflect the power of training size to predict the phenotypes in future years. **Figures 3 and 4** show the results for forward predictions at Pullman and Lind when combined data from the previous years were used to predict the phenotypes. The X-axis represents the year for which predictions were made while training the models on all the previous year’s phenotypic data (**Figure 3 & 4**). We saw a gradual increase in prediction accuracy for all the traits as we kept increasing the training data size, and the same trend was observed for both locations (**Figure 3 & 4**). The highest improvement in prediction accuracy was observed for GPC, FPROT, FASH, and FSDS, owing to the complex nature of these traits and demonstrating the importance of training size. Similar to cross-validation prediction accuracy (**Table 3**), the highest forward prediction accuracy was obtained with machine and deep learning models, especially when the training data size kept increasing (**Figure 3 & 4**). Bayesian models performed worst for all of the traits and at both locations, even when training data size was increased.

Forward predictions in 2019 were, on average, 32% and 29% greater than the forward predictions in 2016 for Pullman and Lind (**Figure 3 & 4**). The highest improvement in forward predictions from 2016 to 2019 was 0.35 to 0.55 for CODI, while the lowest was 0.26 to 0.29 for KWT (**Figure 3**). The highest improvement was seen for MLP and CNN, demonstrating as the size of training data increases, deep learning models result in the highest improvement in prediction accuracy. Furthermore, cross-validation prediction accuracies were, on average, 34% and 32% more than forward prediction in 2019 for Pullman and Lind (**Table 3, Figure 3 & 4**), suggesting that cross-validation scenarios over-inflate prediction accuracies.

### Across location predictions

Across location predictions were performed where data from Lind was used to train the model for predicting performances in Pullman and vice versa. Owing to all the Bayesian model’s worst performance and computational burden in cross-validation and forward predictions, these models were eliminated for the across location predictions. **Figure 5 and Table 4** showed the prediction accuracy for all fourteen end-use quality traits when predictions were made for 2019_Pullman by models training on the whole Lind dataset and vice versa. The across location prediction accuracies were, on average, 16% and 47% less than forward and cross-validation prediction accuracies, demonstrating the importance of inclusion of genotype by environment interaction components into the GS models for across location and environment predictions.

**Table 4.** Genomic selection across environment prediction accuracies for fourteen end-use quality traits evaluated with four different models. 2019_Pullan_Lind denotes the scenario where 2019_Pullman was predicted using datasets from Lind as the training set and vice versa for 2019_Lind_Pullan. The highest accuracy for each trait is bolded under different model scenarios.

| Location | Trait | RRBLUP | RF | MLP | CNN |
| --- | --- | --- | --- | --- | --- |
| 2019_Pullman_Lind | FYELD | 0.41 | 0.48 | **0.50** | 0.46 |
|  | BKYELD | 0.31 | 0.38 | 0.38 | **0.40** |
|  | MSCOR | 0.27 | **0.30** | **0.30** | **0.30** |
|  | TWT | 0.32 | 0.37 | **0.38** | **0.38** |
|  | GPC | 0.25 | 0.30 | 0.31 | **0.33** |
|  | KHRD | 0.32 | 0.37 | 0.36 | **0.38** |
|  | KWT | 0.34 | **0.37** | 0.36 | 0.36 |
|  | KSIZE | 0.34 | 0.38 | 0.38 | **0.40** |
|  | CODI | 0.40 | 0.45 | **0.46** | **0.46** |
|  | FPROT | 0.35 | 0.40 | 0.40 | **0.41** |
|  | FASH | 0.40 | 0.41 | 0.41 | **0.42** |
|  | FSV | 0.27 | 0.36 | **0.39** | 0.36 |
|  | FSDS | 0.36 | **0.44** | 0.43 | 0.41 |
|  | FSRW | 0.36 | 0.39 | 0.41 | **0.42** |
| 2019_Lind_Pullman | FYELD | 0.43 | 0.47 | **0.50** | 0.49 |
|  | BKYELD | 0.31 | 0.40 | **0.41** | 0.40 |
|  | MSCOR | 0.28 | 0.29 | **0.31** | **0.31** |
|  | TWT | 0.31 | 0.36 | 0.35 | **0.37** |
|  | GPC | 0.27 | 0.30 | 0.28 | **0.31** |
|  | KHRD | 0.33 | 0.33 | **0.38** | 0.37 |
|  | KWT | 0.34 | 0.37 | **0.38** | 0.37 |
|  | KSIZE | 0.35 | 0.39 | **0.40** | **0.40** |
|  | CODI | 0.42 | 0.44 | **0.46** | **0.46** |
|  | FPROT | 0.34 | **0.42** | **0.42** | 0.40 |
|  | FASH | 0.41 | **0.42** | **0.42** | 0.40 |
|  | FSV | 0.30 | 0.38 | 0.38 | **0.42** |
|  | FSDS | 0.38 | **0.41** | 0.40 | 0.40 |
|  | FSRW | 0.37 | 0.41 | 0.41 | **0.43** |
| Average |  | **0.34** | **0.38** | **0.39** | **0.39** |

All the abbreviation are previously abbreviated in the text and Table 2.

Deep learning models performed best for across location prediction compared to RRBLUP and machine learning models (**Table 4 & Figure 5**). The highest prediction accuracy was 0.50 for FYELD with a MLP model for predicting 2019_Pullman (**Table 4**). The lowest prediction accuracies were for MSCOR, GPC, and FSV with the RRBLUP model for predicting 2019_Pullman (**Table 4**). Out of the four models used, twelve end-use quality traits were best predicted by deep learning models under the 2019_Pullman scenario, while RF performed best for the remaining two traits (**Table 4**). In 2019_Lind predictions, the highest accuracy again 0.50 for FYELD with the MLP model, and lowest was for GPC and MSCOR with the RRBLUP model. Similar to 2019_Pullman, deep learning models performed best for eleven out of the fourteen traits evaluated in 2019_Lind.

## Discussion

Selection for end-use quality traits is often more difficult to conduct compared to grain yield, disease resistance, and agronomic performance, due to the cost, labor, and seed quantity requirements (Chhabra et al. 2021). Phenotyping for quality traits is usually delayed until later generations, resulting in creating small population sizes with unbalanced datasets (Battenfield et al. 2016). This study explored the potential of GS, especially machine and deep learning models, for predicting fourteen different end-use quality traits using five years (2015-19) of phenotyping data from a winter wheat breeding program. The prediction accuracy in this study varied from 0.27-0.81, demonstrating the potential of its implementation in the breeding program. We observed that forward and across location prediction accuracies could be increased using deep and machine learning models without accounting for genotype by environment interaction, environment covariates, and kernel matrices in traditional GS models. Furthermore, QTLs or major genes controlling quality traits are typically already fixed in the particular market class or breeding programs; hence, GS is the best substitute for marker-assisted selection by exploring different combinations of QTL to achieve the best variety (Lorenz 2013).

The broad-sense heritability of end-use quality traits evaluated varied from 0.56 to 0.93, with the majority of them having a value above 0.80. Similar heritability values were obtained by Michel et al. (2018), Jernigan et al. (2017), and Kristensen et al. (2019) for different baking and flour yield parameters of winter wheat. These intermediate to high heritability estimates suggested that most of the variation in these traits is genetic and less affected by environment and genotype by environment interactions (Tsai et al. 2020). Therefore, GS is the best option for predicting these traits due to capturing most of the additive genetic variation by the models, as observed in this study, due to intermediate to high prediction accuracy for different quality traits. We observed that only a few grain and flour assessments traits were correlated. These low correlations among most end-use quality traits strengthen the fact that no single quality parameter can assist in final variety selection, but that many are needed (Souza et al. 2002). Only three end-use quality traits, namely, GPC, FPROT and FSV, had intermediate heritability values, which were also reported in previous studies due to their complex and polygenic inheritance nature (Hayes et al. 2017; Sandhu et al. 2021c). Similarly, comparatively low prediction accuracies obtained from these traits validated the fact for inclusion of genotype by environment interaction or environmental covariates for their prediction (Monteverde et al. 2019).

Cross-validation prediction accuracies were, on average, 34% and 32% higher than forward prediction in 2019 for Pullman and Lind. The higher cross-validation prediction accuracies compared to forward and across location prediction suggests the importance of including bigger training sets, genotype by environment interactions, and environment covariates for exploiting the maximum variation to make predictions (Gouy et al. 2013). Higher accuracies obtained in cross-validation showed that most of those values are over-inflated, and attention is required before making any final decision about those large values to adopt GS in the breeding program (Crossa et al. 2014). Cross-validation approaches included training and testing sets from the same environment, thus accounting for environmental variation in prediction. Moreover, most of the lines evaluated in breeding programs are usually closely related or full sibs and confound cross-validation approaches, where full sibs might be in the same training or testing group, causing inflation in prediction accuracies (Rutkoski et al. 2015). The relationship between individuals in the training and testing set profoundly affects model performance, with a closer relationship resulting in higher accuracy. Forward and across location prediction are the best method for studying the importance of GS implementation in the breeding program (Habier et al. 2013; Fiedler et al. 2017).

Continuous increments in forward prediction accuracy with all nine models demonstrated the importance of a large training population and more environments for training the GS model (Yao et al. 2018). He et al. (2016) and Battenfield et al. (2016) observed an increase in forward prediction in spring wheat end-use quality traits. Similarly, Meuwissen et al. (2016) suggested updating the GS model with a large training population every cycle to increase prediction accuracy. They observed a rise in genetic gain for fertility, longevity, milk production, and other traits in cows by following this. Deep learning models saw the greatest improvement in forward prediction accuracy by including more training data and new environments, supporting the importance of big data for their best performance (Cuevas et al. 2019). Furthermore, across location predictions were superior by using deep learning models. This could be attributed to capturing genetic, environmental, and genotype by environment interaction components by these models without explicit programming (Montesinos-López et al. 2019c). Across location prediction can be further improved by including genotype by environment interactions or environment covariates like weather or soil parameters into the GS models to make across location and environment selections (Jarquín et al. 2014; Monteverde et al. 2019).

We observed differences in model prediction accuracies under all scenarios evaluated in this study, where machine and deep learning models performed superior to Bayesian and RRBLUP models. This difference in model performance is attributed to the different genetic architecture of each trait, dependent upon the heritability and number of QTLs controlling that trait (Plavšin et al. 2021). Similar results were obtained by various other studies showing the superiority of machine learning models over conventional Bayesian models in wheat (Gianola et al. 2006; Montesinos-López et al. 2019a; Merrick and Carter 2021). Hu et al. (2019) showed that random forest performed superior to the Bayesian and RRBLUP for predicting thousand kernel weight, grain protein content, and sedimentation volume in wheat under forward prediction scenario, further strengthening our findings that machine and deep learning models should be explored for such conditions. Furthermore, we observed that highly heritable traits in this study have higher prediction accuracy than moderately heritable traits, suggesting that in addition to genetic architecture, the heritability of a trait also plays an important role in final prediction accuracy (Huang et al. 2016; Hayes et al. 2017).

Machine and deep learning models performed better than all Bayesian and RRBLUP models under cross-validation, forward, and across location predictions. The higher prediction accuracy observed due to deep and machine learning models is attributed to their flexibility in deciphering complex interactions between responses and predictors to capture different trends present in the datasets compared to only additive variation in conventional GS models (Montesinos-López et al. 2021). Deep and machine learning models explore the whole feature space during model training using different sets of neurons, activation function, and various other hyperparameters to identify the best pattern for giving the best prediction scenario compared to Bayesian models that include a pre-selected prior distribution for final predictions. Furthermore, most of the traits were predicted best by different deep and machine learning due to their respective genetic architecture of each trait. Some studies in wheat reported that all models give the same prediction accuracy irrespective of the model used while others strengthen the superiority of different models for different traits (Heslot et al. 2015; Schmidt et al. 2016). Ma et al. (2018) and Montesinos-López et al. (2019) also obtained similar results to predict quantitative traits in wheat and suggested that deep learning models should be explored due to their better prediction accuracies.

It is believed that machine and deep learning models should be used on very large training datasets, which is often not possible for end-use quality traits which are evaluated at later stages of the breeding process. However, this and other studies have shown that even small dataset can give equivalent or superior performance to the traditional parametric GS models (Ma et al. 2018; Pook et al. 2020; Sandhu et al. 2021a). Moreover, Bellot et al. (2018) have used a training set of 100k individuals and showed no advantage of deep learning models over the conventional GS models. Pérez□Rodríguez et al. (2020) and Liu et al. (2019) showed the superiority of different deep learning algorithms over conventional GS models using population sizes of 268 wheat and 4294 soybean lines. These results provide evidence that training datasets play a minor role in prediction compared to the genetic architecture of the trait, but the importance of large population sizes in GS models still can’t be undermined. The main issue with using a small dataset for deep learning models is overfitting, resulting in the model’s failure to learn the exact pattern from the dataset (Montesinos-López et al. 2021). Herein, we used regularization and dropout functions to remove a certain number of neurons during model training to avoid the overfitting problem (Srivastava et al. 2014; Lecun et al. 2015).

### Conclusion

We assessed the potential of machine and deep learning genomic selection models for predicting fourteen different end-use quality traits at two locations in a soft white winter wheat breeding program. Different cross-validation, forward, and across location prediction scenarios were tried for comparing different models and utilization of this approach in the breeding program. Owing to limited seed availability, time constraint, and associated cost, phenotyping for quality traits is delayed to later generations. However, the higher accuracy of prediction models observed in this study suggest that selections can be performed earlier in the breeding process. Machine and deep learning models performed better than Bayesian and RRBLUP genomic selection models and can be adopted for use in plant breeding programs, regardless of dataset sizes. Furthermore, the increase in forward prediction accuracy with the addition of more lines in the training set concluded that genomic selection models should be updated every year for the best prediciton accuracy. Overall, this and previous studies showed the benefit of implementing genomic selection with machine and deep learning models for different complex traits in large scale breeding programs using collected phenotypic data from previous years.

## Acknowledgements

We would like to thank Kerry Balow, Adrienne Burke, Gary Shelton, and Kyall Hagemeyer for assisting in population development, genotyping, and field plot maintenance.

## Authors contribution

Conceptualization: KSS, MA, & AHC; Writing original draft: KSS; Data analysis: KSS; Genotyping curation and filtration: MA; Review and editing: KSS, MA, CM, & AHC; Resources: CM & AHC; Supervision and Funding: AHC. All authors approved the final document for submission.

